# Under Cover of Darkness: Refuge from Artificial Light at Night may Mitigate Risks to Stranded Seabirds

**DOI:** 10.1101/2024.08.12.607643

**Authors:** Taylor M. Brown, Kaitlyn Baker, Sabina I. Wilhelm, Gary Burness

## Abstract

Artificial light at night is an anthropogenic pollutant that has wide-ranging effects on wildlife. Fledgling seabirds of the order Procellariiformes exhibit phototaxis toward artificial lights on their first flights from the nest, causing them to become grounded in human settlements where they are subject to increased risk of predation. Limited evidence suggests that certain light types may be less attractive than others, yet there is also evidence for an aversion to light under certain circumstances. We investigated differences in phototactic behaviour, activity level, and shelter-seeking behaviour of grounded *Hydrobates leucorhous* (Leach’s storm-petrel) fledglings exposed to artificial light in three experiments: a Y-maze choice experiment, an open field test, and a modified open field test with a hide box provided (“Safe Haven test”). When provided with combinations of different light types in the Y-maze, storm-petrels typically remained stationary in the darkest part of the apparatus and exhibited no response toward one light type over another. This was consistent with results from the open field test, in which individuals were less active in darkness than when exposed to two out of three light conditions (Warm White Light-Emitting Diode and High Pressure Sodium). More than half of birds entered the hide box in light conditions, compared to none in the dark. Considered together, our results indicate Leach’s storm-petrel fledglings exhibit photophobic behaviour after stranding, which may be part of a behavioural strategy to avoid predation. Further, we demonstrate the utility of providing hide boxes to protect stranded seabird fledglings in locations where lighting cannot be eliminated or where rescue efforts may be limited in spatial or temporal coverage. However, hide boxes would have limited utility in dark locations. Hide boxes constitute a novel mitigation measure that merits future testing for its ability to reduce stranding-induced mortality, especially in imperiled procellariiform species.

**LAY SUMMARY:**

- Globally, fledglings of many seabird species become stranded in towns and at industrial sites near their breeding colonies
- Strandings are thought to be caused by attraction to lights, although some species appear to avoid light
- We tested behavioural responses and activity levels of Leach’s storm-petrel fledglings toward several different light types
- When presented with different types of light in a choice experiment, most birds remained stationary
- Fledglings were more active in warm-hued lights than in darkness
- Birds sought shelter when it was provided in light conditions, but not when provided in darkness
- Our results suggest that rescue programs targeting Leach’s storm-petrels should include searches in dark, hidden locations where fledglings are likely to hide, and support the use of small shelters to protect stranded seabirds when lights cannot be turned off

## INTRODUCTION

Nightscapes across our planet are increasingly subject to light pollution from anthropogenic sources. Further, the widespread conversion to energy-efficient Light-Emitting Diode (LED) lighting is expected to worsen ecological impacts through a variety of mechanisms, including an increase in the usage of short light wavelengths that are especially disruptive to wildlife (Pawson and Bader 2014, Davies and Smyth 2018, Jägerbrand and Spoelstra 2023). The presence of artificial light at night (ALAN) can alter the physiology and behaviour of a growing list of animals (Sanders et al. 2021, Yang et al. 2024). Especially in nocturnal animals, artificial light can act as a supranormal visual stimulus that induces phototactic behaviour (movement either toward or away from light; Rich and Longcore 2006). Positive phototaxis has been observed in many animal groups including insects, amphibians, fish, sea turtles, migratory songbirds, and seabirds, among others (Rich and Longcore 2006, Rodríguez et al. 2017a, Jägerbrand and Spoelstra 2023). Phototaxis toward ALAN sources can result in mortality through a variety of mechanisms including trauma from collision with buildings, vehicles, and other structures; dehydration or desiccation after stranding in unfamiliar locations; and predation or poaching of disoriented individuals (Witherington 1997, Rich and Longcore 2006, Rodríguez et al. 2017a, Van Doren et al. 2021).

Seabirds of the order Procellariiformes are one of the most imperiled groups of birds, with anthropogenic threats including fisheries bycatch, plastic ingestion, climate change, and predation by introduced predators (Dias et al. 2019). In addition, fledglings of burrow-nesting and largely nocturnal species of procellariiform, such as storm-petrels and shearwaters, are susceptible to stranding in developed coastal areas and at offshore industrial sites after disorientation by, and sometimes collision with, lighted structures during their first flights from the nest (sometimes termed “fallout”; Montevecchi 2006, Burke et al. 2012, Rodríguez et al. 2017a, Gjerdrum et al. 2021). Fledgling mortality following stranding can be high without rescue intervention (Rodríguez et al. 2017a, Burt et al. 2024). Even where rescue programs are implemented, many stranded fledglings are not found as they tend to hide in concealed locations (Reed et al. 1985, Rodríguez et al. 2017a). Although procellariiforms are generally K-selected and their populations are therefore relatively robust against juvenile mortality, light-induced fallout has had a significant negative effect on population growth in several species (Ainley et al. 2001, Fontaine et al. 2011, Griesemer and Holmes 2011, Simons 1984) and adults can also be affected (Rodríguez et al. 2017a, Burt et al. 2024).

Increasing experimental evidence suggests that reductions in ALAN near colonies of burrow-nesting, night-fledging seabirds can decrease stranding numbers (Reed et al. 1985, Miles et al. 2010, Rodríguez et al. 2014, Brown et al. 2024, Burt et al. 2024). At finer scales and in ground-based contexts, however, behavioural patterns are more equivocal and vary among species. For example, in the charadriiform *Synthliboramphus antiquus* (Ancient Murrelet), chicks orient more often toward reflected light sources than toward darkness, yet because they nest in remote areas and go to sea before they can fly are not prone to fallout (Gaston et al. 1988). Stranding-prone *Fratercula arctica* (Atlantic Puffin; Order Charadriiformes) fledglings similarly orient toward light over darkness in a Y-maze (Brown et al. 2024), but stranded *Calonectris diomedea* (Cory’s shearwater; Order Procellariiformes) fledglings consistently move toward darkness in a similar experiment (Atchoi et al. 2024). The latter result may indicate a context-dependent phototactic response in which behaviour toward ALAN on a broad scale (especially in flight) differs from fine-scale responses (especially on the ground, post-stranding). Combined with observational and tracking data of fledgling seabirds as they encounter ALAN (e.g., Rodríguez et al. 2022), it has been suggested that procellariiform fledglings may be attracted to artificial lights from a distance but disoriented by them at close range (Atchoi et al. 2024).

In instances where total or near-total suppression of ALAN is not possible (e.g., industrial sites with nocturnal operations), other mitigation measures are needed to reduce fallout. The shielding of light from radiating upward toward birds flying overhead has demonstrated effectiveness (Reed et al. 1985, Urmston et al. 2022). Implementation of potentially “less attractive” light types also has potential. Limited evidence suggests that long wavelength-dominant (i.e., “redder”) lights (such as sodium vapour types) with low correlated colour temperature (CCT; a relatively crude but simple and widely-used metric of the perceived colour of a nominal white light source; Durmus 2022) may be less attractive to fledgling procellariiforms and other wildlife than those with high-CCT, short wavelength-dominant (i.e., “bluer”) spectra (such as many LEDs; Rodríguez et al. 2017b, Longcore et al. 2018). Increased attraction toward blue-violet wavelengths may be explained by an increased sensitivity to these wavelengths by the optical systems of the affected taxa (Reed 1986, Bowmaker et al. 1997, Hart 2004, Pawson and Bader 2014, Davies and Smyth 2018).

Contrary results, however, have been found in Y-maze experiments with Cory’s Shearwaters, which preferentially oriented toward red over blue light (Atchoi et al. 2024); and with Atlantic Puffins, which showed no differences in response between blue and orange light or between low- and high-CCT LED light (Brown et al. 2024). Interestingly, however, activity levels of puffin fledglings did appear to vary by light type: birds were more active when exposed to High Pressure Sodium (HPS) light compared to LED light (Brown et al. 2024). These results imply effects of light type on behaviour beyond simple phototaxis: for example, varying activity levels following stranding under different light types could have implications for detection probability of individuals by rescue programs. To date, no similar research has been conducted on activity levels under various light types of any procellariiform, despite this group’s overwhelming representation in the list of light-affected seabirds.

*Hydrobates leucorhous* (Leach’s storm-petrels; Order Procellariiformes, Family Hydrobatidae) are one of many seabird species affected by stranding. They are a small (approximately 60 grams at fledging), long-lived (20-30+ years), colonial, burrow-nesting species that breeds on islands in the North Atlantic and North Pacific oceans (Pollet et al. 2021). The largest colony of this species is on Baccalieu Island in Newfoundland and Labrador, Canada, with an estimated breeding population of 1.95 million pairs (down from an estimated 3.4-5.1 million pairs in 1984; Wilhelm et al. 2020). Juveniles fledge on average two hours after sunset in September through November (Collins et al. 2023), going directly to the open ocean without parental guidance (Pollet et al. 2021). At this time, some fledglings become stranded both inland and at offshore oil production and other industrial facilities (Miles et al. 2010, Gjerdrum et al. 2021, Wilhelm et al. 2021, Burt et al. 2023, 2024). Leach’s storm-petrels fledge throughout the lunar cycle but stranding increases when moon illumination is low (Miles et al. 2010, Collins et al. 2023, Burt et al. 2024), potentially implying that artificial light has a greater attractive or disorienting effect on birds during nights with lower natural light levels (Rodríguez et al. 2023).

On the Avalon Peninsula of Newfoundland and Labrador, hundreds to thousands of Leach’s storm-petrels are found stranded each year and many likely remain unfound, especially given the large geographic area over which strandings occur (Gjerdrum et al. 2021, Wilhelm et al. 2021, Burt et al. 2023). High predation on grounded individuals by *Larus argentatus* (Herring Gulls), *Vulpes* spp. (foxes), *Felis catus* (feral cats), and other predators at “hotspot” stranding sites (Burt et al. 2024) as well as high mortality of stranded procellariiforms from other causes even with the use of rescue programs (Rodríguez et al. 2017a) imply the need for additional mitigation measures where lights cannot be extinguished and where rescuers cannot be constantly present. For example, the targeted placement and monitoring of protective hiding places at stranding “hotspots” may be beneficial to reduce predation (Burt et al. 2024), as procellariiforms often seek shelter in concealed locations after grounding (Reed et al. 1985, Rodríguez et al. 2017a).

Our objectives were to: 1) measure phototactic behavioural responses of grounded Leach’s storm-petrel fledglings (hereafter “storm-petrels”) to artificial lights; and 2) measure their activity under various lighting conditions. Our working hypothesis across both objectives was that storm-petrels are attracted to artificial light, and their levels of phototactic behaviour and general activity will vary across light spectra. We used a Y-maze to measure phototactic behaviour (objective 1) and expected that fledglings would exhibit positive phototaxis by preferentially orienting toward light over darkness when both were provided as options. We further expected that when storm-petrels were exposed to two different artificial light types, they would orient more often toward those with “cooler” hues (e.g., Cool White LED; Blue LED) than to those with “warmer” hues (e.g., High Pressure Sodium; Warm White LED; Orange LED). We addressed objective 2 using an open field test: although we expected to observe differences in activity levels among the light treatments, particularly when compared with darkness, we could not predict directionality *a priori*. Lastly, and based on anecdotal observations of grounded storm-petrels seeking shelter in rodent bait stations (Burt et al. 2024), we leveraged the results from objectives 1 and 2 to test a third objective: whether different lighting conditions would influence an individual’s propensity to seek shelter in a hide box (termed a “Safe Haven” box). This latter objective has the potential to inform local-scale rescue efforts and conservation measures.

## METHODS

### Study Sites and Species

Experiments were conducted with stranded storm-petrels in October 2021 and 2023 at the Quinlan Brothers seafood processing plant in Bay de Verde (48.085°N, 52.895°W), Newfoundland and Labrador, Canada. Hundreds of storm-petrels strand at the seafood processing plant on an annual basis (Wilhelm et al. 2021, Burt et al. 2023, 2024), most of which likely originate from Baccalieu Island, approximately 7 km to the northeast (Figure 1). We conducted experiments in two sheds on the wharf between 21:00 and 04:30 on 2-11 October 2021 and 3-8 October 2023.

**Figure 1.**
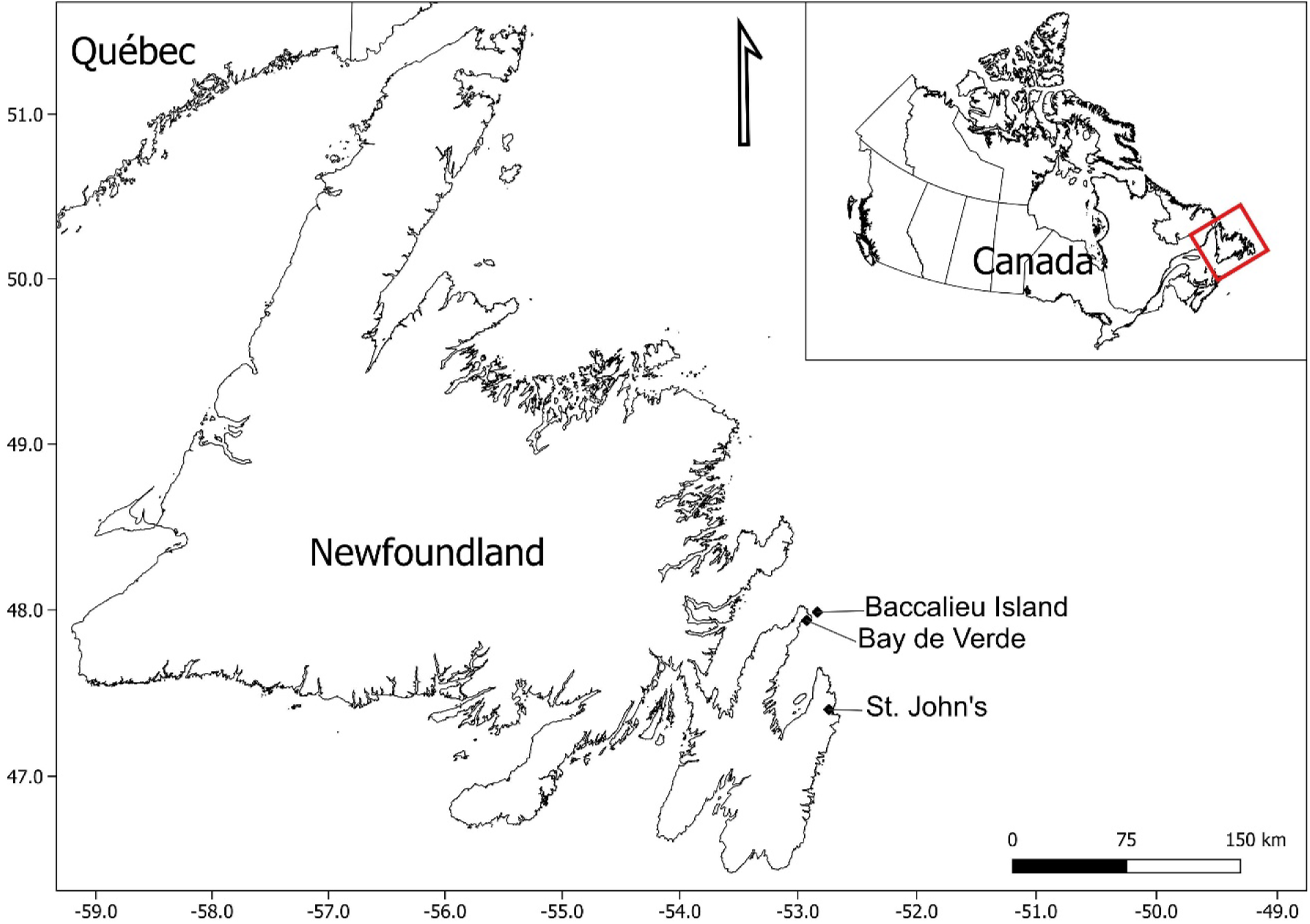
Leach’s storm-petrel colony location (Baccalieu Island) and study location (Bay de Verde) with reference to the capital city (St. John’s) of the province of Newfoundland and Labrador, Canada.

### Animal Collection, Housing, and Release

We collected stranded storm-petrels during opportunistic walking surveys (single “laps”) around the processing plant from approximately 20:00 to 24:00. We conducted experiments throughout the night between or at the same time as walking surveys, depending on the number of field personnel available (range 1-5). Although it is possible that stranded storm-petrel fledglings do not represent a truly random sample of the entire population, it is this population in which we are most interested since it appears to be most affected by ALAN and the population from which we would expect the greatest response to our experimental light stimuli. When individuals were found, we placed them in cardboard pet carriers, approximately 47 cm long by 25 cm wide by 30 cm tall and lined with disposable absorbent multipurpose pads (up to 25 individuals per carrier). We kept the carriers containing rescued birds in darkness in a third shed until experimentation and/or release.

Prior to experimentation (described below), we removed each storm-petrel from the group housing and placed it individually in a cardboard box (30 cm × 20 cm × 25 cm) outdoors in a sheltered location for a minimum 10-minute rest period. Following the experiment testing phototactic behaviour (in 2021), we replaced the bird in its cardboard box and provided another rest period of minimum 10 minutes before participating in the activity level experiment. In 2023, the Safe Haven test involved two trials in close succession: storm-petrels used in this experiment rested for a minimum of 10 minutes before the first trial, but for only approximately 5 minutes between the first and second trial. In both years, once an individual had completed its trials, we placed it in a cardboard pet carrier with other birds that had already completed experimental trials, until release.

### Choice Experiment to Measure Phototactic Behaviour

To measure phototactic responses of storm-petrels to various light spectra we used a Y-maze, following Brown et al. (2024). Briefly, the apparatus consisted of a dark, metal “acclimation box” (34 × 15 × 15 cm) which opened into a plastic “main box” (40 × 55 × 22 cm) via a vertical sliding door. From the main box, a pair of corrugated plastic pipes led to one of two plastic choice boxes (36 × 27 × 18 cm). Each choice box had a 3 cm-diameter round hole in the end, through which our experimental lights could shine into each choice arm. We used two infrared baby monitor cameras (VTech, VM5262-2; British Columbia, Canada) to view the doorway between each choice arm and its corresponding choice box, through small holes in the walls of the choice boxes, allowing us to detect when an individual had made a response.

We used three different types of lights in combination with various filters to produce four stimulus combinations (materials and spectra of all light-filter combinations used can be found in Supplementary Material Table S1). The lights used were: High Pressure Sodium (HPS) with a CCT of 2100 K; Warm White LED with a CCT of 2700 K; and Cool White LED with a CCT of 5000 K (see Supplemental Material). We randomly selected 76 stranded storm-petrel fledglings and divided them amongst four light stimulus combinations. The combinations (with number of trials each, in brackets) were: HPS vs Darkness (*n* = 19); HPS vs Warm White LED (*n* = 19); Warm White LED vs Cool White LED (*n* = 17); and Blue LED vs Orange LED (*n* = 21). The blue and orange LED light options were created by adding to the Cool and Warm White LED options filters made of coloured acrylic and glass (Table S1). The various light spectra are found in Figure S2. We tested only one combination each night, selected randomly without replacement until all four had been tested over the course of four nights; they were then selected again in random order (all light combinations tested over two non-sequential nights).

Before each trial, we placed the light stimuli in their positions and turned them on. We brought the storm-petrel into the choice experiment shed, removed it from its box, and placed it in the acclimation box of the Y-maze. We set a timer for two minutes, after which we removed the inner sliding door, and the bird was allowed 13 minutes to make a “response”. If the bird did not voluntarily exit the acclimation box within 5 minutes after removal of the inner door, it was gently prodded with the experimenter’s hand until it entered the main box (*n* = 53 out of 76 storm-petrels, or 70%, were prodded). We considered the bird to have completed a response when it reached the end of the choice arm, and any part of the body was visible on the baby monitor screen (even if it did not fully enter the choice box); otherwise, the trial outcome was deemed “No Response”. Note that not all prodded birds made responses; prodding was merely used in an attempt to elicit a response. The bird was removed from the apparatus after making a response or after 15 minutes, whichever came first, and returned to its cardboard box in preparation for the activity level experiment. In most cases of “No Response,” storm-petrels sat still in the apparatus, but “No Response” also includes the few individuals that audibly wandered around inside the apparatus without ever becoming visible on the monitor screen. Between trials we cleaned the entirety of the Y-maze with 70% isopropyl alcohol and a disposable cloth and allowed it to dry completely. We then swapped the two lights between the two choice boxes to account for any potential side bias on part of the birds.

### Measuring Activity Levels Under Various Light Spectra

We used an open field test to measure differences in activity level of fledgling storm-petrels under various artificial light spectra, as described in Brown et al. (2024). Briefly, the 1-m³ cube-shaped open field test arena was constructed from polycarbonate panels, with the floor left unattached for easy removal. We created a 20 cm × 20 cm grid composed of 25 squares (i.e., 5 squares × 5 squares) on the floor, cut an access door into the bottom edge of one wall of the arena for easy insertion and removal of storm-petrels from the apparatus, and cut two small holes in the ceiling: one for the light source and one for the camera (Activeon, CCA10W, Activeon Inc.; San Diego, CA, USA). We placed an opaque blackout curtain over the whole arena to prevent any ambient light from entering.

The same 76 storm-petrel fledglings used in the Y-maze were divided into four treatment groups for trials in the open field test, where each individual was exposed to darkness or one of three light spectra. Two individuals were removed from the final dataset due to camera malfunction, resulting in a final sample size of *n* = 74. Treatments were: Darkness (*n* = 21); HPS (*n* = 18); Warm White (2700K) LED (*n* = 16); and Cool White (5000K) LED (*n* = 19). We did not measure activity in response to blue or orange light to ensure adequate sample sizes in the light treatments that most closely resemble actual light spectra used in human settlements and at industrial sites. Median illuminance at floor level in the arena was 23 – 26 lux and did not differ significantly across the three light treatments (Brown et al. 2024). Each night, we tested two light types in the open field test (chosen randomly), and they were never the same as the two light types used in the Y-maze on the same night. From the remaining light options, the initial treatment each night was chosen randomly, followed by the second treatment, and these were alternated for the rest of the night.

Prior to each open field trial, we set up and turned on the appropriate light (except in the Dark treatment, in which the light was in place but not turned on) and started a video recording (with audio); unfortunately, a thermal imaging camera was not available for this experiment. We placed the storm-petrel just inside the access door of the arena and started an electronic timer for 10 minutes, after which time the video and audio recording were stopped. The storm-petrel was removed from the arena and replaced in its box. The floor of the arena was not cleaned between trials unless a storm-petrel defecated during its trial, in which case the affected area was cleaned with 70% isopropyl alcohol and allowed to dry before the next trial. We completed as many trials as possible each night (median 11 trials; range 3-13 trials per night).

We scored open field test videos in a random order. We used behavioural scoring software JWatcher (Version 1.0, Macquarie University and University of California, Los Angeles) to code two levels of behaviour for each 10-minute trial: “mobile”, which included walking, running, or flying, and “immobile” whenever birds were stationary. The software automatically calculated the total amount of time each storm-petrel spent mobile and immobile. For storm-petrels in the “Dark” treatment group, only audio data were discernible. Storm-petrels were audible when they walked on the polycarbonate floor of the arena, which allowed us to hear and score their movement. As such, rather than viewing the file, we instead listened to the audio file and coded “mobile” behaviours when we heard the bird walking or flying, and “immobile” when we heard the bird stop moving. Scratching (clawing at the arena floor with the feet) could also occasionally be heard but was not scored as “mobile” behaviour on its own. To determine whether the time spent mobile in the Dark treatment could be accurately scored by listening to the audio alone, we selected a random subset of 10 videos from the three lighted conditions (HPS, Warm White LED, Cool White LED) that had already been scored visually. We listened to all 10 of these videos in random order and scored them audially in the same way that we scored the Dark videos. A single observer (KB) did all visual and audial scoring. We then performed a linear regression and used the resulting equation (*y* = 0.7*x* + 56) to convert the Dark treatment data for use in later analyses (see Supplemental Material).

### Safe Haven Test

We used the same polycarbonate arena for the Safe Haven test as we used for the open field test; however, we removed the roof and its overlaying curtain. Inside the arena we placed an empty black plastic rodent bait station (approximately 29 cm × 25 cm × 15 cm; Bell Laboratories Protecta Evo Express; Wisconsin, USA) which we hereafter refer to as the “Safe Haven box” (Figure 2). The Safe Haven box has two small openings (approximately 6 cm in diameter) through which storm-petrels can enter; one on each of the two short sides. We placed the Safe Haven box against the middle of the wall opposite the arena’s access door, with its two openings oriented along the wall. Inside the Safe Haven box, a ramp angles upward from each opening until dropping off to the interior floor of the box, such that after an individual goes over the ramp, it has difficulty escaping the box.

**Figure 2.**
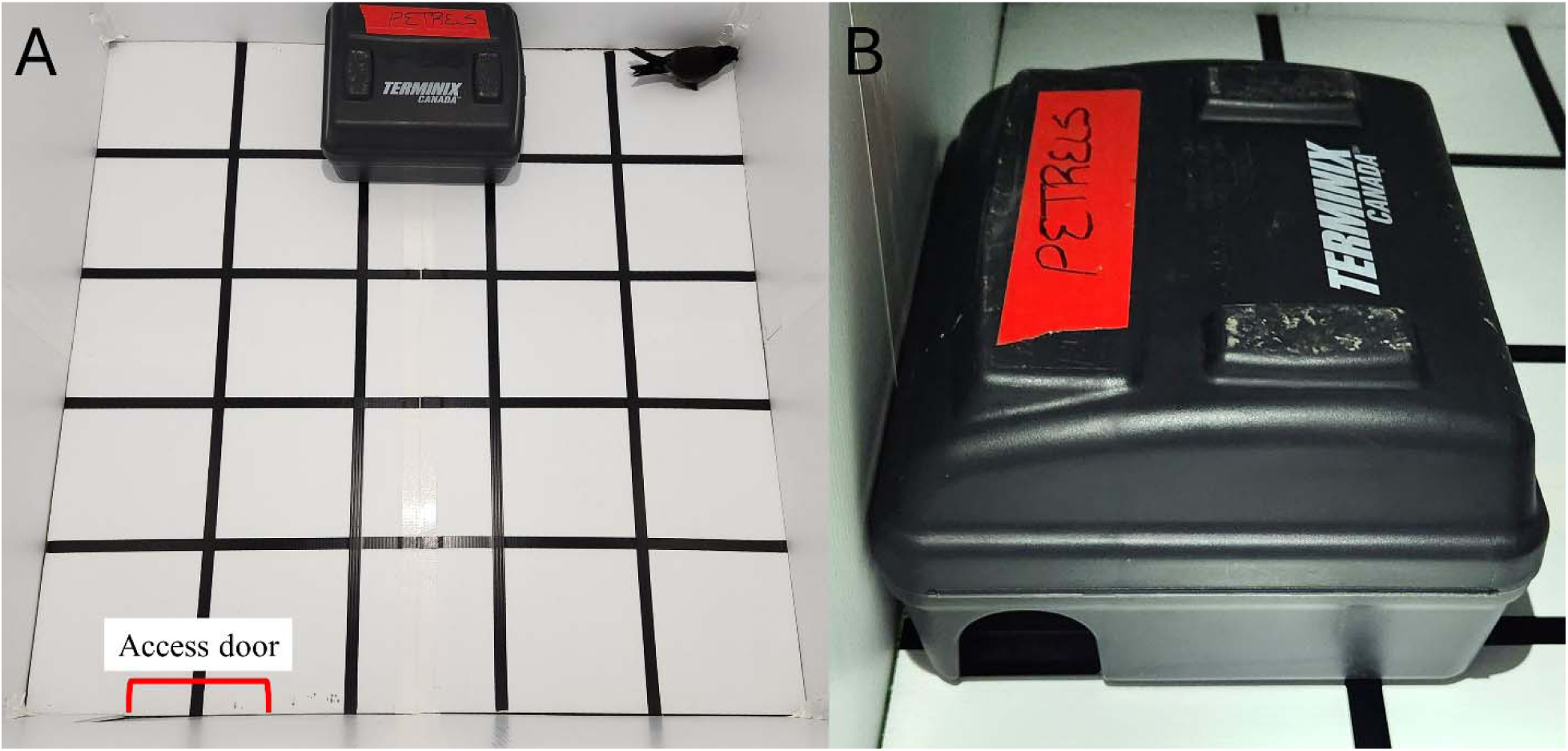
A) Floor layout of the Safe Haven test arena, including Safe Haven box and storm-petrel. Note the access door at the bottom of the picture, through which storm-petrels were placed inside the arena. Each floor grid square is 20 cm × 20 cm. B) Orientation of the Safe Haven box such that the openings were both against the wall of the arena.

This experiment consisted of two treatments: Light and Dark. The light treatment used a Cool White 5000 K LED bulb as in the previous experiments, hung approximately 160 cm above the centre of the arena floor. In the Dark treatment, all light sources in the experiment arena were extinguished and windows in the shed were covered. To record the behaviour of individuals in the arena we used a thermal imaging camera (Teledyne FLIR C5; Oregon, USA) affixed to a tripod and positioned such that it could record the entire arena floor.

Between 3-8 October 2023, we tested 14 storm-petrels in a paired design, such that each individual experienced one Light trial and one Dark trial (Light trials, *n* = 14; Dark trials, *n* = 14). We chose randomly whether the first bird experienced Light or Dark first and alternated the first treatment experienced by each bird thereafter, to obtain a roughly equal number of individuals that experienced Light first (*n* = 6) versus Dark first (*n* = 8). In each trial, we started the video camera, placed each storm-petrel just inside the access door of the arena, and started a timer. At the end of each 10-minute trial, we noted whether the storm-petrel was inside the Safe Haven box and, if it was not, we later used the thermal imaging video footage to verify that the bird had not entered the Safe Haven box during the experiment (note that there were zero occurrences of this). We then removed the storm-petrel from the arena (or from the Safe Haven box, when applicable), replaced it in its cardboard holding box, and allowed it to rest for 5 minutes until the arena was ready for the second trial. The floor of the arena was not cleaned between trials unless a storm-petrel defecated during its trial. If a storm-petrel entered the Safe Haven box, we cleaned the entire Safe Haven box with 70% isopropyl alcohol and allowed it to dry before the next trial.

### Statistical Analysis

All analyses were performed in R (v. 4.3.0; R Core Team 2023). We report results to a significance level of *α* = 0.05.

### Choice experiment to measure phototactic behaviour

To test whether storm-petrels displayed phototactic responses toward certain light spectra over others, we first excluded individuals who exhibited “No Response”. We used the remaining data to conduct exact two-tailed binomial tests (package “*stats*”, v. 4.3.0; R Core Team 2023) on the numbers of storm-petrels in each stimulus combination that responded to each of the two provided light options. We tested against a hypothesized proportion of 0.5. We report 95% confidence intervals of the estimated true proportions.

### Measuring activity levels under various light spectra

We conducted a Levene Test (package “*car*”, v. 3.1-2; Fox and Weisberg 2019) to test for homogeneity of variance in time spent mobile across treatments, and a Shapiro-Wilk test (package “*stats*”, v. 4.3.0; R Core Team 2023) on the residuals of a standard ANOVA of time spent mobile across treatments to test for normality. Variance differed significantly across treatments (*F* = 7.5594; df = 3 and 70; *p* < 0.001) but the there was no evidence for deviation from normality (*W* = 0.984, *p* = 0.5). We therefore used a Welch’s one-way ANOVA (which does not assume equal variances) to determine if the time spent mobile differed significantly among treatment groups, followed by a pairwise post-hoc Games-Howell test since it does not assume homogeneity of variance or equal sample sizes (package “*rstatix*”, v. 0.7.2; Kassambara 2023).

### Safe haven test

To test whether individual storm-petrel fledglings differed in their probability to enter the Safe Haven box depending on the light treatment used (Light or Dark), we conducted a classical version of McNemar’s chi-squared test for paired data (Pembury Smith and Ruxton 2020).

## RESULTS

### Choice Experiment to Measure Phototactic Behaviour

Sixty-three percent, i.e. 48 of the 76 storm-petrels displayed “No Response” (Table 1) and were excluded from subsequent binomial tests to evaluate light response. Of the remaining 28 individuals that responded to one of the two provided light options, 43%, i.e. 12 were prodded out of the acclimation box into the main box after 5 minutes of inactivity, while 57%, i.e. 16 voluntarily left the acclimation box before making their response (Table 1). Focusing on the 28 storm-petrels that made a response, we found no significant response toward any light type in our stimulus combinations (i.e., light type 1 vs. light type 2; Table 2). Among individuals that made a response, there was also no statistical evidence for side bias, as 15 storm-petrels chose Box 1 and 13 storm-petrels chose Box 2 (binomial test *p* = 0.9; 95% CI of estimated true proportion of Box 1: 0.34-0.72).

**Table 1.** Contingency table comparing the number of storm-petrels that required prodding out of the acclimation box after 5 minutes of inactivity with the number of individuals that eventually made a response, or not, in the Y-maze choice experiment. Also shown are totals in each category and total experimental sample size (*n* = 76).

|  |  | Prodded |  |  |
| --- | --- | --- | --- | --- |
|  |  | Yes | No | Total |
| Responded | Yes | 12 | 16 | 28 |
|  | No | 41 | 7 | 48 |
| Total |  | 53 | 23 | **76** |

**Table 2.** Number of storm-petrels that responded to each of the two light options in each stimulus combination (columns 2 and 4) or exhibited no response for one light type over the other (“No Response”). We also report the 95% confidence intervals of the true proportion of “Light type 1” as a reference, and the *p*-value of the exact two-tailed binomial test on the quantities that responded to each light option.

| Light type 1 | Number | Light type 2 | Number | 95% CI | $p$-value | No response |
| --- | --- | --- | --- | --- | --- | --- |
| HPS | 3 | Dark | 6 | 0.07-0.70 | 0.5 | 10 |
| HPS | 5 | Warm White | 3 | 0.24-0.91 | 0.7 | 11 |
| Warm White | 4 | Cool White | 4 | 0.16-0.84 | 1.0 | 9 |
| Blue | 2 | Orange | 1 | 0.09-0.99 | 1.0 | 18 |

### Measuring Activity Levels Under Various Light Spectra

The time that storm-petrels spent mobile differed significantly among light treatments (Welch’s one-way ANOVA *F*_3,30.28_ = 9.306, *p* < 0.001). Specifically, we found that individuals spent significantly less time mobile in Darkness (mean time spent mobile = 84 seconds) than in Warm White LED light (mean time spent mobile = 210 seconds) and in HPS light (mean time spent mobile = 172 seconds; *p* < 0.01 and *p* < 0.05, respectively; Figure 3). None of the other light treatments differed significantly from each other in time spent mobile. Note that there are no scores of time spent mobile lower than 56 seconds in the Dark group. This was because of the conversion applied to scores of time spent mobile in Darkness using a linear equation with intercept *y* = 56 (see Supplemental Material). Although this adds a degree of uncertainty to some scores in the Dark group (some individuals were almost certainly active for < 56 seconds), it results in a more conservative approach to estimating differences in time spent mobile across groups.

**Figure 3.**
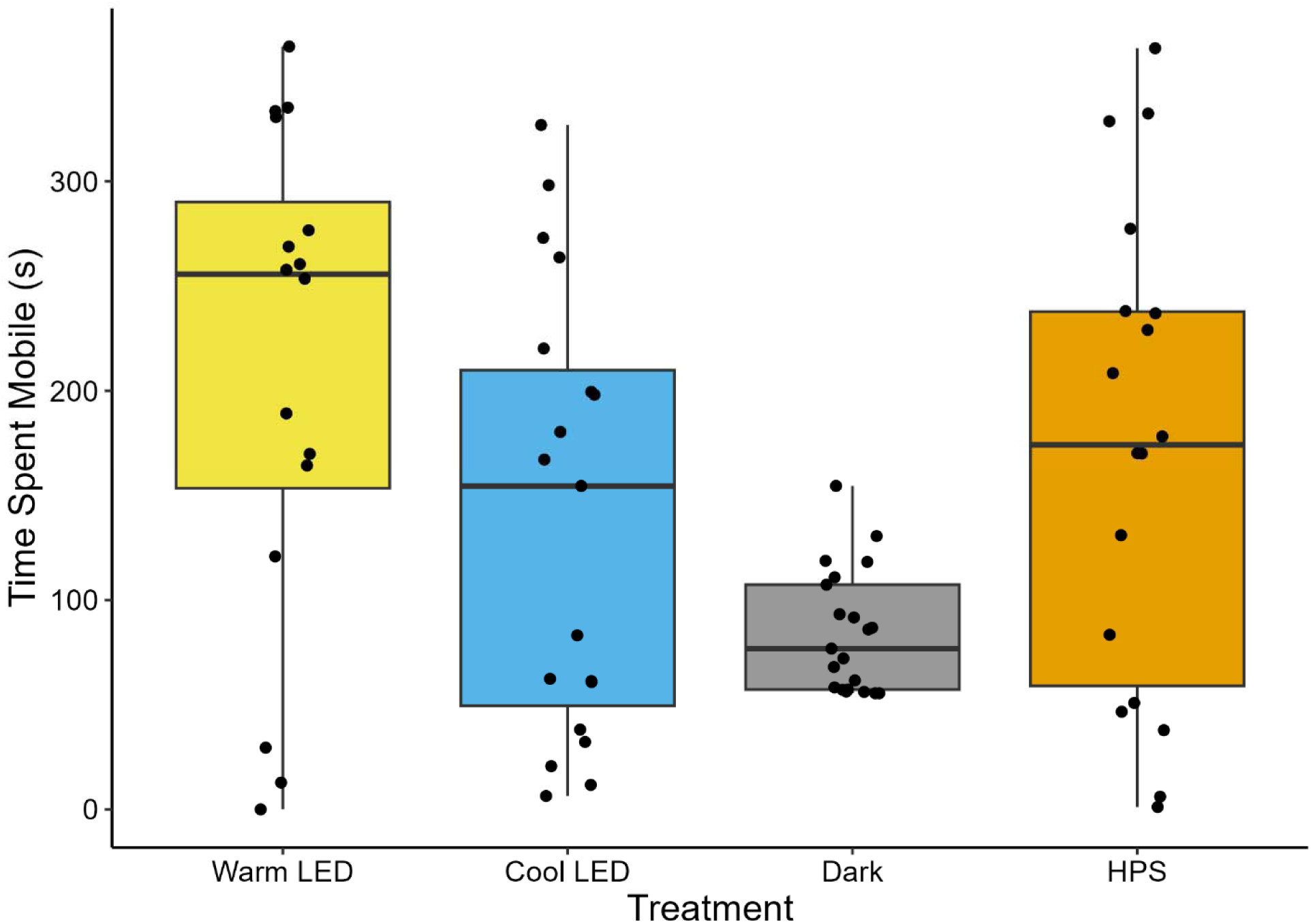
Fledgling storm-petrels were significantly less mobile in the Dark than when exposed to Warm White LED or High Pressure Sodium (HPS). There were no other significant differences. Figure shows a boxplot of time spent mobile (in seconds) by individuals exposed to one of four light treatment groups, with overlaid individual data points. Note that, because of the conversion applied to Dark data (see text), data points of birds that truly spent zero or near-zero time mobile have been artificially inflated and statistical comparisons are therefore more conservative.

### Safe Haven Test

Storm-petrels were significantly more likely to enter the Safe Haven box in light conditions than in the dark (classical McNemar’s test χ^2^(1) = 6.125, *p* < 0.05). In the dark, none of the 14 individuals entered the Safe Haven box. In contrast, when exposed to the Light treatment, 8 of the same 14 storm-petrels entered the Safe Haven box while 6 did not enter.

## DISCUSSION

We expected that storm-petrels would display evidence of positive phototaxis when tested in a laboratory setting. In contrast, most fledglings either displayed no response to light stimuli or displayed evidence of negative phototaxis (light avoidance). Across our three experiments, birds tended to remain stationary in darkness, while in light conditions they exhibited increased movement. In the Y-maze, storm-petrels exhibited little response to the light options provided and most remained relatively immobile in both the dark acclimation box and, following prodding, in the (also relatively dark) main box of the Y-maze apparatus. Further, although there were no significant responses displayed in any stimulus combination, numerically more birds moved toward darkness in the HPS vs Darkness group (three and six, respectively). The apparent photophobic behaviour of storm-petrels in the Y-maze choice experiment was consistent with the lower activity levels they exhibited in darkness in our open field test, and with the increased probability of storm-petrels to enter hide boxes in light compared to dark conditions. Specifically, our Safe Haven Test results provide evidence that the purpose of increased activity levels in light conditions was to seek out a dark place to hide.

Our findings are consistent with those of Atchoi et al. (2023, 2024), who found evidence for photophobic behaviour in fallout-susceptible Cory’s shearwater chicks and fledglings (but not adults) when they were exposed to light in controlled experimental apparatuses. Anecdotally, seabird fledglings are sometimes found in dark and concealed locations after grounding, implying negative phototaxis (Reed et al. 1985, Rodrigues et al. 2012, Burt et al. 2024). Our results also concur with the photophobia exhibited by adult burrow-nesting procellariiforms (including Leach’s storm-petrels) at their colonies in the form of reduced attendance levels and later arrival when light is at naturally high levels (Watanuki 1986, Riou and Hamer 2008, Silva et al. 2011) or artificially introduced (Syposz et al. 2021, Austad et al. 2023).

Instead of attraction toward ALAN being the sole mechanism driving procellariiform strandings, lights may attract birds from a distance and then cause disorientation at close range, as evidenced in part by the highly tortuous flights of fledglings as they navigate over brightly-illuminated areas in comparison with dark areas (Rodríguez et al. 2022). Negative phototaxis by fledgling procellariiforms in Y-maze apparatuses (where light stimuli are presented at very close range) has been interpreted as birds attempting to avoid the disorientating stimulus (Atchoi et al. 2024). Alternatively, but not mutually exclusively, once birds are grounded and more vulnerable to predation, they may become negatively phototactic and seek the safety of darkness. Our experiment cannot distinguish these competing hypotheses; future studies might seek to test them by introducing artificial visual, auditory, or even olfactory predator cues into an open field or Y-maze experiment to determine if photophobic behaviour is increased in the presence of a perceived predator.

Our initial (incorrect) prediction that storm-petrel fledglings would be positively phototactic in our Y-maze choice experiment was based on the observations that fledglings of this and other burrow-nesting seabird species are found grounded each year in anthropogenically developed areas, presumably due to light attraction (Rodríguez et al. 2017a, Wilhelm et al. 2021, Burt et al. 2023), and that fallout increases with increased ALAN (e.g., Miles et al. 2010, Brown et al. 2024, Burt et al. 2024). In contrast to fledgling procellariiforms, fledglings of burrow-nesting charadriiforms display positive phototaxis on the ground (specifically those in the family Alcidae; Gaston et al. 1988, Brown et al. 2024). It is unclear why the two groups differ despite their shared tendencies to develop in burrows, fledge nocturnally and become grounded by ALAN. It has been suggested that in burrow-nesting procellariiforms the visual system is not fully developed at the time of fledging (itself due to a lack of light exposure in the burrow throughout development; Mitkus et al. 2018), which may contribute to the disorientation they experience in response to ALAN encountered on their first flights from the nest (Atchoi et al. 2020); presumably, the same would be true of burrow-nesting charadriiforms. In general, burrow-nesting charadriiforms also share with procellariiforms the same rod and cone photoreceptors and their associated spectral sensitivities (Reed 1986, Ebrey and Koutalos 2001, Ödeen and Håstad 2003). However, still little is known about variation in ocular anatomy, physiology, and development across stranding-prone seabird species and how they may contribute to observed differences in phototactic behaviour.

When provided with different light spectra in a Y-maze, we expected storm-petrels to preferentially respond to short wavelength-dominant spectra (e.g., Cool White LED; Blue LED) over those that are long wavelength-dominant (e.g., Warm White LED; Orange LED), based on increased numbers of stranded individuals in the presence of short wavelength-dominant light (Salamolard et al. 2001 as cited in Minatchy 2004, Rodríguez et al. 2017b). Our current results demonstrating no differences in response by storm-petrels to different light types agree with those of our previous Y-maze experiment with Atlantic Puffin fledglings (Brown et al. 2024), but differ from those of Cory’s shearwaters, in which both fledglings and adults oriented more toward red than blue light (Atchoi et al. 2024). This behaviour was speculated to be an adaptation for Cory’s shearwaters to avoid disorientation (Atchoi et al. 2024), since procellariiforms may be especially sensitive to and disoriented by blue-rich light (Hart 2004, Rodríguez et al. 2017b, Syposz et al. 2021). In comparison, few storm-petrel fledglings responded to either blue or orange light in our Y-maze, with 86% (18 out of 21) exhibiting “No Response” in that combination. This suggests either a tendency to remain in the dark, or an aversion to both types of spectra in storm-petrels. Alternatively, storm-petrels may have perceived differences in brightness among our provided light options which led to unquantified effects on the behavioural responses (or lack thereof) they exhibited in our experiments. We calibrated most of our light stimuli such that the amount of light irradiated at the peak wavelength of each spectrum was relatively equal (Figure S2), but this does not mean that all spectra were perceived as equally bright. Nevertheless, we did this because the calibration of spectra based on perceived “brightness” is difficult without a complete understanding of the spectral sensitivities and ocular physiology in the species of interest.

In our open field test, storm-petrels spent significantly less time mobile in darkness than when exposed to either Warm White LED or HPS light. This suggests that storm-petrels are particularly stimulated by spectra rich in long-wavelength (i.e., “reddish”), but not short wavelength (i.e., “bluish”), light. We did not find any differences in activity levels among the three light treatments (HPS, Warm White LED, or Cool White LED). These results are in direct contrast with those of Atlantic Puffin fledglings, which are highly active in darkness as well as HPS light and less active in both Warm and Cool White LED light (Brown et al. 2024). To our knowledge, activity levels in response to artificial light have not been quantified in any other seabird species. Low activity levels of storm-petrels in darkness could imply either a calming or startling effect from the lack of light; the mechanism underlying this behavioural pattern remains unexplained. Nonetheless, these results hold implications for detection probability of stranded birds by rescuers whereby presumably, mobile birds are more conspicuous than stationary birds, all else being equal.

When fledging birds become grounded, they are vulnerable to increased predation risk (Rodríguez et al. 2017a, Burt et al. 2024). To address our third objective, we tested the propensity of storm-petrel fledglings to enter a provided dark hiding place under light versus darkness. We found that more than half of birds entered in light conditions, compared to none in the dark. This clearly demonstrates the value of placing Safe Haven boxes (in this case empty rodent bait stations) in lit areas to provide stranded storm-petrels with protection until rescue, as had previously been suggested by Burt et al. (2024). Additionally, anecdotal preliminary trials revealed that birds never entered the Safe Haven box when its doors were oriented away from the wall of the open field apparatus, so this needs to be considered when deploying them. During initial deployments of Safe Haven boxes at our study site (beginning April 2021; Burt et al. 2024), we noted up to 11 individuals sheltering inside a single box in a single night (TMB and SIW, pers. obs.), demonstrating their effectiveness in a field setting. Safe Haven boxes therefore provide an effective and inexpensive means to protect at least a portion of grounded seabirds and potentially further reduce stranding-related mortality in stranding “hotspots”, a tactic especially helpful in trying to further conserve already declining and threatened populations affected by ALAN.

## Conclusion

We found substantial evidence for photophobic behaviour in fledgling storm-petrels that have been grounded in areas illuminated by ALAN. Importantly, we have also demonstrated that this behaviour tends to be exhibited regardless of the spectrum of light to which birds are exposed. Our study, which focused on small-scale behavioural responses to light in a ground-based context, does not predict phototactic behaviour of birds in flight or how it may relate to stranding. Further, the storm-petrels used in our experiments were already grounded and may have encountered any number of stressors prior to capture that could have influenced their behaviour. It must therefore be remembered that seabird umwelt remains largely a mystery and that behavioural responses in artificial versus natural settings will inevitably differ to some degree. Our findings nonetheless have implications for seabird rescue programs: procellariiforms that become grounded in dark areas will likely remain there, as evidenced by their low activity levels and lack of hiding response in experimentally dark conditions. This is in contrast to procellariiform fledglings grounded in light areas, which likely exhibit high activity levels and seek out dark hiding places (as also observed anecdotally in previous studies). Given that individuals become stranded in large numbers in lighted areas and rescue efforts are typically focused there, our results emphasize the need to also search more concealed dark locations where stranded birds may hide. Our results also confirm the utility of an additional mitigation measure not previously tested in the global effort to reduce mortality of seabirds grounded by artificial light at night. The extinguishing of ALAN is not always possible, especially in industrial settings where human safety during nocturnal operations is a priority. The deployment of Safe Haven boxes in illuminated stranding “hotspots” takes advantage of the negatively phototactic behaviour displayed by stranded fledglings. This will assist in the safe rescue of fledglings and provide an additional measure to help conserve seabird populations affected by ALAN.

## LITERATURE CITED

Ainley DG, R Podolsky, L Deforest, G Spencer, and N Nur (2001). The status and population trends of the Newell’s shearwater on Kaua’i: Insights from modeling. Studies in Avian Biology 22:108–123.

Atchoi E, M Mitkus, and A Rodríguez (2020). Is seabird light-induced mortality explained by the visual system development? Conservation Science and Practice 2:e195. 10.1111/csp2.195

Atchoi E, M Mitkus, P Vitta, B Machado, M Rocha, M Juliano, J Bried, and A Rodríguez (2023). Ontogenetic exposure to light influences seabird vulnerability to light pollution. Journal of Experimental Biology 226:jeb245126. 10.1242/jeb.245126

Atchoi E, M Mitkus, B Machado, V Medeiros, S Garcia, M Juliano, J Bried, and A Rodríguez (2024). Do seabirds dream of artificial lights? Understanding light preferences of Procellariiformes. Journal of Experimental Biology 227:jeb247665. 10.1242/jeb.247665

Austad M, S Oppel, J Crymble, HR Greetham, D Sahin, P Lago, BJ Metzger, and P Quillfeldt (2023). The effects of temporally distinct light pollution from ships on nocturnal colony attendance in a threatened seabird. Journal of Ornithology 164:527–536. 10.1007/s10336-023-02045-z

Bowmaker JK, LA Heath, SE Wilkie, and DM Hunt (1997). Visual pigments and oil droplets from six classes of photoreceptor in the retinas of birds. Vision Research 37:2183–2194. 10.1016/S0042-6989(97)00026-6

Brown TM, SI Wilhelm, AD Slepkov, K Baker, GF Mastromonaco, and G Burness (2024). Navigating the night: effects of artificial light on the behaviour of Atlantic Puffin fledglings. Animal Behaviour 218:135–148. 10.1016/j.anbehav.2024.09.008

Burke CM, WA Montevecchi, and FK Wiese (2012). Inadequate environmental monitoring around offshore oil and gas platforms on the Grand Bank of Eastern Canada: are risks to marine birds known? Journal of Environmental Management 104:121–126. 10.1016/j.jenvman.2012.02.012

Burt TV, SM Collins, and WA Montevecchi (2023). Social media reports inform the spatio-temporal distribution of Leach’s storm-petrel strandings across the island of Newfoundland. Northeastern Naturalist 30:151–160. 10.1656/045.030.0204

Burt TV, SM Collins, S Green, PB Doiron, SI Wilhelm, and WA Montevecchi (2024). Reduction of coastal lighting decreases seabird strandings. PLoS ONE 19(6): e0295098. 10.1371/journal.pone.0295098

Collins SM, A Hedd, WA Montevecchi, T Burt, DR Wilson, and DA Fifield (2023). Leach’s storm-petrels fledge on the full moon and throughout the lunar cycle. Biology Letters 19: 20230290. 10.1098/rsbl.2023.0290

Davies TW, and T Smyth (2018). Why artificial light at night should be a focus for global change research in the 21^st^ century. Global Change Biology 24:872–882. 10.1111/gcb.13927

Dias MP, R Martin, EJ Pearmain, IJ Burfield, C Small, RA Phillips, O Yates, B Lascelles, PG Borboroglu, and JP Croxall (2019). Threats to seabirds: a global assessment. Biological Conservation 237:525–537. 10.1016/j.biocon.2019.06.033

Durmus D (2022). Correlated color temperature: use and limitations. Lighting Research & Technology 54:363–375. 10.1177/14771535211034330

Ebrey T, and Y Koutalos (2001). Vertebrate photoreceptors. Progress in Retinal and Eye Research 20(1):49–94. 10.1016/S1350-9462(00)00014-8

Fontaine R, O Gimenez, and J Bried (2011). The impact of introduced predators, light-induced mortality of fledglings and poaching on the dynamics of the Cory’s shearwater (*Calonectris diomedea*) population from the Azores, northeastern subtropical Atlantic. Biological Conservation 144:1998–2011.

Fox J, and S Weisberg (2019). An R Companion to Applied Regression, Third edition. Sage, Thousand Oaks CA. <https://socialsciences.mcmaster.ca/jfox/Books/Companion/>.

Gaston AJ, IL Jones, DG Noble, and SA Smith (1988). Orientation of ancient murrelet, Synthliboramphus antiquus, chicks during their passage from the burrow to the sea. Animal Behaviour 36:300–303. 10.1016/S0003-3472(88)80277-X

Gjerdrum C, RA Ronconi, KL Turner, and TE Hamer (2021). Bird strandings and bright lights at coastal and offshore industrial sites in Atlantic Canada. Avian Conservation and Ecology 16(1):22. 10.5751/ACE-01860-160122

Griesemer AM, and ND Holmes (2011). *Newell’s shearwater population modeling for habitat conservation plan and recovery planning*. Technical report No. 176. The Hawai’i-Pacific Islands cooperative ecosystem studies Unit & Pacific Cooperative Studies Unit. Honolulu, HI: University of Hawai’i, 68 pp.

Hart NS (2004). Microspectrophotometry of visual pigments and oil droplets in a marine bird, the wedge-tailed shearwater *Puffinus pacificus*: topographic variations in photoreceptor spectral characteristics. Journal of Experimental Biology 207:1229–1240. 10.1242/jeb.00857

Jägerbrand AK, and K Spoelstra (2023). Effects of anthropogenic light on species and ecosystems. Science 380:1125–1130. 10.1126/science.adg3173

Kassambara A (2023). rstatix: Pipe-Friendly Framework for Basic Statistical Tests. R package version 0.7.2, <https://CRAN.R-project.org/package=rstatix>.

Longcore T, A Rodríguez, B Witherington, JF Penniman, L Herf, and M Herf (2018). Rapid assessment of lamp spectrum to quantify ecological effects of light at night. Journal of Experimental Zoology Part A 329:511–521. 10.1002/jez.2184

Miles W, S Money, R Luxmoore, and RW Furness (2010). Effects of artificial lights and moonlight on petrels at St Kilda. Bird Study 57:244–251. 10.1080/00063651003605064

Minatchy N (2004). Stratégie de réduction de la mortalité des pétrels induite par les éclairages publics. Specialized diploma dissertation, University of Réunion, Réunion.

Mitkus M, GA Nevitt, and A Kelber (2018). Development of the visual system in a burrow-nesting seabird: Leach’s storm petrel. Brain, Behavior and Evolution 91:4–16. 10.1159/000484080

Montevecchi WA (2006). Influences of artificial light on marine birds. In Ecological Consequences of Artificial Night Lighting (pp. 94–113). Washington, DC: Island Press.

Ödeen A, and O Håstad (2003). Complex distribution of avian color vision systems revealed by sequencing the SWS1 opsin from total DNA. Molecular Biology and Evolution 20(6):855–861. 10.1093/molbev/msg108

Pawson SM, and MK-F Bader (2014). LED lighting increases the ecological impact of light pollution irrespective of color temperature. Ecological Applications 24:1561–1568. 10.1890/14-0468.1

Pembury Smith MQR, and GD Ruxton (2020). Effective use of the McNemar test. Behavioral Ecology and Sociobiology 74:133. 10.1007/s00265-020-02916-y

Pollet IL, AL Bond, A Hedd, CE Huntington, RG Butler, and R Mauck (2021). Leach’s Storm-Petrel (*Hydrobates leucorhous*), version 1.1. In Birds of the World (Editor not available). Cornell Lab of Ornithology, Ithaca, NY, USA.

R Core Team (2023). R: A language and environment for statistical computing. R Foundation for Statistical Computing, Vienna, Austria. URL https://www.R-project.org/.

Reed JR (1986). Seabird vision: spectral sensitivity and light-attraction behavior. PhD thesis, University of Wisconsin – Madison, Wisconsin, USA.

Reed JR, JL Sincock, and JP Hailman (1985). Light attraction in endangered procellariiform birds: reduction by shielding upward radiation. Auk 102(2):377–383.

Rich C, and T Longcore (Eds.). (2006). *Ecological consequences of artificial night lighting*. Island Press.

Riou S, and KC Hamer (2008). Predation risk and reproductive effort: impacts of moonlight on food provisioning and chick growth in Manx shearwaters. Animal Behaviour 76:1743–1748. 10.1016/j.anbehav.2008.08.012

Rodrigues P, C Aubrecht, A Gill, T Longcore, and C Elvidge (2012). Remote sensing to map influence of light pollution on Cory’s shearwater in São Miguel Island, Azores Archipelago. European Journal of Wildlife Research 58:147–155.

Rodríguez A, G Burgan, P Dann, R Jessop, JJ Negro, and A Chiaradia (2014). Fatal attraction of short-tailed shearwaters to artificial lights. PLoS ONE 9(10): e110114. 10.1371/journal.pone.0110114

Rodríguez A, ND Holmes, PG Ryan, K-J Wilson, L Faulquier, Y Murillo, AF Raine, JF Penniman, V Neves, B Rodríguez, JJ Negro, A Chiaradia, P Dann, T Anderson, B Metzger, M Shirai, L Deppe, J Wheeler, P Hodum, C Gouveia, V Carmo, GP Carreira, L Delgado-Alburqueque, C Guerra-Correa, F-X Couzi, M Travers, and M Le Corre (2017a). Seabird mortality induced by land-based artificial lights. Conservation Biology 31(5):986–1001.

Rodríguez A, P Dann, and A Chiaradia (2017b). Reducing light-induced mortality of seabirds: High pressure sodium lights decrease the fatal attraction of shearwaters. Journal for Nature Conservation 39:68–72.

Rodríguez A, B Rodríguez, Y Acosta, and JJ Negro (2022). Tracking flights to investigate seabird mortality induced by artificial lights. Frontiers in Ecology and Evolution 9:786557. 10.3389/fevo.2021.786557

Rodríguez A, E Atchoi, B Rodríguez, T Pipa, M Le Corre, and DG Ainley (2023). Moonlight diminishes seabird attraction to artificial light. Conservation Science and Practice 5:e13014. 10.1111/csp2.13014

Salamolard M, F-X Couzi, M Pellerin, and T Ghestemme (2001). Etude complémentaire concernant la protection du Puffin de Baillon dans le cadre de la Route des Tamarins. Rapp. SEOR/DDE-Réunion. 30 pp. + Annexes 4 pp.

Sanders D, E Frago, R Kehoe, C Patterson, and KJ Gaston (2021). A meta-analysis of biological impacts of artificial light at night. Nature Ecology & Evolution 5:74–81. 10.1038/s41559-020-01322-x

Silva MC, JP Granadeiro, PD Boersma, and I Strange (2011). Effects of predation risk on the nocturnal activity budgets of thin-billed prions *Pachyptila belcheri* on New Island, Falkland Islands. Polar Biology 34:421–429. 10.1007/s00300-010-0897-6

Simons TR (1984). A population model of the endangered Hawaiian dark-rumped petrel. Journal of Wildlife Management 48:1065–1076.

Syposz M, O Padget, J Willis, BM Van Doren, N Gillies, AL Fayet, MJ Wood, A Alejo, and T Guilford (2021). Avoidance of different durations, colours and intensities of artificial light by adult seabirds. Scientific Reports 11:18941. 10.1038/s41598-021-97986-x

Urmston J, KD Hyrenbach, and K Swindle (2022). Quantifying Wedge-tailed shearwater (*Ardenna pacifica*) fallout after changes in highway lighting on southeast O’ahu, Hawai’i. PLoS ONE 17(3): e0265832. 10.1371/journal.pone.0265832

Van Doren BM, DE Willard, M Hennen, KG Horton, EF Stuber, D Sheldon, AH Sivakumar, J Wang, A Farnsworth, and BM Winger (2021). Drivers of fatal bird collisions in an urban center. PNAS 118: e2101666118. 10.1073/pnas.2101666118

Watanuki Y (1986). Moonlight avoidance behavior in Leach’s storm-petrels as a defense against Slaty-backed gulls. The Auk 103:14–22.

Wilhelm SI, A Hedd, GJ Robertson, J Mailhiot, PM Regular, PC Ryan, and RD Elliot (2020). The world’s largest breeding colony of Leach’s Storm-petrel *Hydrobates leucorhous* has declined. Bird Conservation International 30:40–57. 10.1017/S0959270919000248

Wilhelm SI, SM Dooley, EP Corbett, MG Fitzsimmons, PC Ryan, and GJ Robertson (2021). Effects of land-based light pollution on two species of burrow-nesting seabirds in Newfoundland and Labrador, Canada. Avian Conservation & Ecology 16(1):12. 10.5751/ACE-01809-160112

Witherington BE (1997). The problem of photopollution for sea turtles and other nocturnal animals. In JR Clemmons & R Buchholz (Eds.), Behavioral approaches to conservation in the wild (pp. 303–328). Cambridge University Press.

Yang Y, Q Liu, C Pan, J Chen, B Xu, K Liu, J Pan, M Lagisz, and S Nakagawa (2024). Species sensitivities to artificial light at night: a phylogenetically controlled multilevel meta-analysis on melatonin suppression. Ecology Letters 27: e14387. 10.1111/ele.14387

