## Supplemental Material for "Under Cover of Darkness: Refuge from Artificial Light at Night may Mitigate Risks to Stranded Seabirds"

**METHODS**

**Statistical Analysis**

**Measuring activity levels under various light spectra.**

To determine if scoring videos using auditory information may have artificially increased or decreased our estimates of time spent mobile in darkness (in comparison to scoring the light videos, which was done visually), we conducted a simple linear regression of time spent mobile scored visually (*y*) as a function of time spent mobile scored audially (*x*) of the 10 selected lighted videos that were scored both visually and using auditory information (package “*stats*”, v. 4.3.0; R Core Team 2023). The regression equation was statistically significant (*r*^2^ = 0.823; *F*(1, 8) = 37.12; *p* < 0.001), and auditory scores predicted visual scores (β = 0.6980; *p* < 0.001; Figure S1). We converted the “Dark” data using the regression equation and used these “converted” values in subsequent statistical comparisons with the unmodified visual scores from the other treatment groups. We decided to score only “Dark” videos using auditory information (rather than scoring all videos this way) because: 1) we reasoned that visual scoring is both more accurate (i.e., truer to the correct value) and more precise (i.e., repeatable) than auditory scoring, and is therefore preferable when three of our four treatments can be scored visually; 2) the noise associated with the High Pressure Sodium light’s ballast fan would have made auditory scoring of those videos difficult and therefore introduce another source of scoring bias; and 3) extra care was taken during the Dark treatment specifically to ensure that noises similar in quality to those of storm-petrel footsteps were reduced to an absolute minimum to increase the accuracy of those auditory scores.

**Supplementary Material Table S1.** Materials used to create each of the six light options used in the choice experiment, including the specific lightbulbs used, whether a neutral-density filter or diffuser were used, and any other materials used.

| **Light option** | **Lightbulb** | **NDF**^a^ | **Diffuser**^b^ | **Other** |
| --- | --- | --- | --- | --- |
| HPS | 2100 K HPS^c^ | yes | yes | none |
| Warm white LED | 2700 K LED^d^ | yes | yes | none |
| Cool white LED | 5000 K LED^e^ | yes | yes | none |
| Blue LED | 5000 K LED^e^ | none | none | ePlastics #2424 dark blue acrylic; cobalt glass |
| Orange LED | 2700 K LED^d^ | yes | yes | ePlastics #2422 amber orange acrylic |
| Darkness | none | none | none | light hole covered |

^a^A variable neutral density filter (NDF) consisted of two pieces of polyvinyl alcohol-iodine polarizing film, overlapping at varying degrees of polarization depending on the light option.
^b^Diffuser consisted of one piece of parchment paper.
^c^High pressure sodium; Light EnerG, 400 Watt
^d^Light-emitting diode; Philips, 5W / 40W-equivalent Soft White LED
^e^Light-emitting diode; Philips, 5W / 40W-equivalent Daylight LED

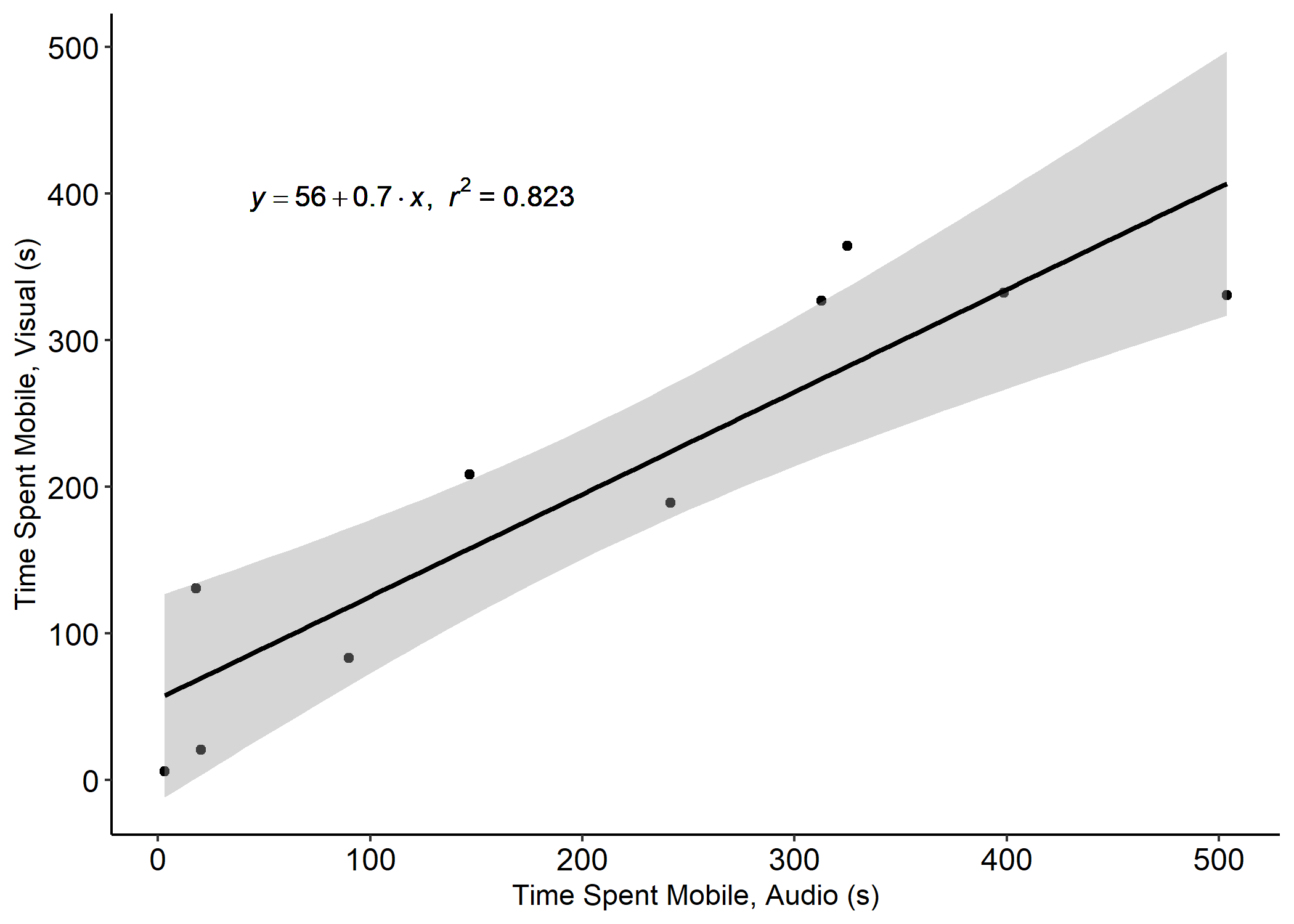

**Supplementary Material Figure S1.** Comparison of time spent mobile by 10 randomly selected Leach’s storm-petrels, each tested in a 600-second trial under either High Pressure Sodium, Warm White LED, or Cool White LED light, as scored by watching the video (y-axis) versus listening to the audio (x-axis). Gray area represents 95% confidence limits.

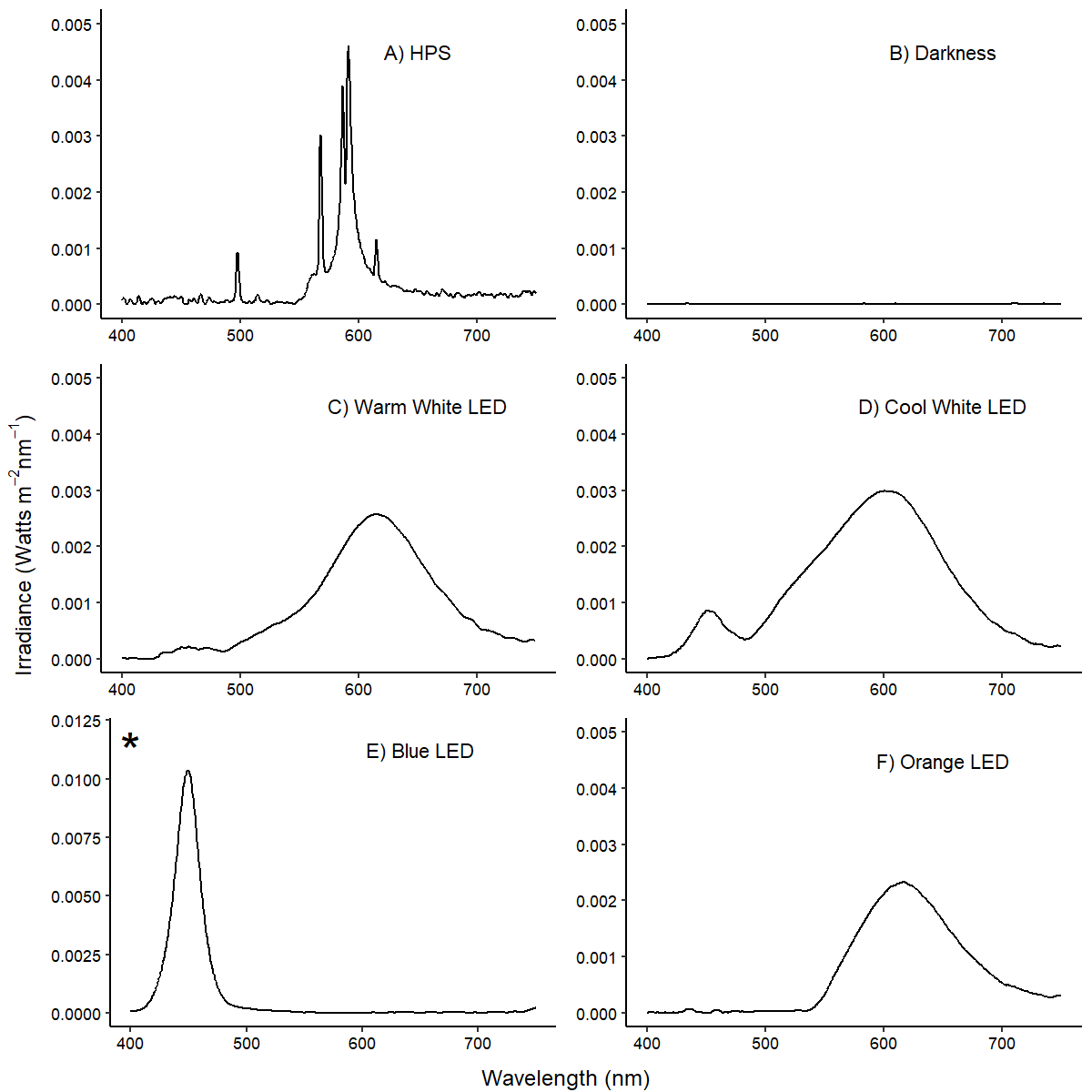

**Supplementary Material Figure S2.** Spectra of all light options used in the choice experiment: A) High pressure sodium; B) Darkness; C) Warm white (2700 K) LED light; D) Cool white (5000 K) LED light; E) Blue LED light; and F) Orange LED light. Irradiance is as measured by a spectrometer with its sensor placed at the entrance of the “choice arm”, inside the “main box” of the Y-maze. * = note the difference in y-axis scale compared to the other spectra.
